# MHCnvex: Likelihood-based model for calling the copy number variations and loss of heterozygosity in MHC class I and II locus

**DOI:** 10.1101/2023.01.29.526131

**Authors:** Noshad Hosseini, Michael B. Mumphrey, Marcin Cieslik

## Abstract

**Motivation:** Accurate detection of copy number variation (CNV) and loss of heterozygosity (LOH) in the major histocompatibility complex (MHC) locus is of great significance to both clinicians and researchers since it has the potential to inform treatment decisions, particularly in the context of immunotherapy. However, due to the high level of polymorphism in this region, calling copy number variations is a challenging task and requires special methodology. To address this challenge, we have developed a tool with a wide range of applicability to call CNV and LOH in the MHC region.

**Results:** To address the challenge mentioned above, we have developed MHCnvex, an algorithm that accurately calls haplotype level CNVs for the genes in the MHC class I and II locus. MHCnvex presents a novel approach based on likelihood models for detecting CNVs at haplotype level. Additionally, this method integrates the MHC locus with other adjacent loci from the short arm of chromosome 6 to enhance the accuracy of the calls. The incorporation of a statistical approach and the examination of the broader chromosome 6 region, rather than just the MHC locus alone, make MHCnvex less vulnerable to local coverage biases (commonly associated with MHC locus). The performance of MHCnvex has been evaluated according to different measures including concordance with MHC flanking regions and changes in the allelic expression of MHC genes due to alteration. MHCnvex has also shown to significantly reduce the variability of calculated coverage for CNV analysis.

**Availability:** Implementation of MHCnvex algorithm is available as an R package at: https://github.com/NoshadHo/MHCnvex

## 1. Introduction

Disrupting Immune surveillance is one of the main routes for malignant cells to survive in the body (*Maleno et al., 2002*). Tumor cells frequently alter their major histocompatibility complex to disrupt cell antigen representing pathway (ATP) (*Gundem et al., 2015*) and therefore escape surveillance by the immune system. One of the main mechanisms by which tumor cells evade the immune system is the Loss of Heterozygosity of MHC genes through copy number alteration. It is critical to understand this mechanism in the context of cancer immunology as it can help to reveal new therapeutic opportunities (*Gundem et al., 2015*). However, High degree of polymorphism in the MHC locus has made it difficult for conventional CNV methods to accurately profile copy numbers for genes in the MHC locus. One of the key challenges arises from the polymorphic nature of MHC region, is to correctly align sequencing reads to the genome. Inaccurate alignment can lead to inaccurate coverage estimates of the locus, thereby causing unreliable CNV calls. Despite the significance of identifying CNVs within the MHC region, there has been a dearth of methodologies developed to address this issue.

In this study, we present MHCnvex, a bioinformatics tool developed as an R package. It utilizes personalized germline references and likelihood modeling to effectively determine MHC CNV and LOH events with high precision. Additionally, MHCnvex integrates MHC locus with adjacent regions on Chromosome 6 to get a full picture of CNV for MHC genes. Inherent noise in sequencing steps makes this integration approach particularly important, as determining copy number variations for short regions of the genome is prone to errors and can lead to local biases in the calculated coverage. Using mentioned methods, MHCnvex can accurately call CNV and LOH for both class-I and class-II MHC genes. To the best of our knowledge, MHCnvex is the only tool that can detect LOH and absolute copy number of genes in both class 1 and class 2, providing a valuable tool for the scientific community. To widen the applicability of the tool, MHCnvex allows for the MHC analysis using WGS, WXS or both if available. In order to increase the utility of MHCnvex in a clinical setting, we have incorporated a “tumor only” method that allows the tool to operate in the absence of a matching normal by using a pool of non-matching normal samples from the same capture kit as tumor. This modification enhances the flexibility of MHCnvex, making it more suitable for use in a clinical setting where matched normal samples may not be readily available.

## 2. Methods

### 2.1 Overview of the algorithm

MHCnvex is an algorithm for copy number analysis of the MHC region, specifically MHC class I and II genes. The MHC locus is the most polymorphic region of the human genome (*Buhler et al, 2011*) and this feature makes it challenging to accurately determine the read coverage and the copy number of MHC genes. To solve this problem, MHCnvex incorporates personalized germline references for the MHC locus which can be inferred by haplotyping tools such as OptiType (*Szolek et al. 2014*) or Hapster (*Mumphrey et al. 2023*). In order to calculate an accurate coverage, MHCnvex utilizes aligned reads to personalized gemline references as input. Furthermore, MHCnvex uses a segmental approach that leverages the fact that CNV events often occur in contiguous segments along the genome. To improve the segmentation accuracy, MHCnvex considers not only the MHC locus itself, but also incorporates adjacent regions into its analysis. This unique approach provides a more comprehensive understanding of the MHC region’s copy-number profile compared to other tools that only focus on the MHC locus alone. Recent years have seen the development of several tools aimed at addressing the challenge of detecting LOH in the MHC region, such as LOHHLA (McGranahan et al., 2017) and DASH (Pyke et al., 2022). However, to the best of our knowledge, none of these methods take into account the potential impact of integrated segmentation of MHC and flanking regions simultaneously on the detection of LOH for the MHC region.

MHCnvex works in the 7 following steps (**Figure 1A**):

1. <u>Coverage calculation:</u> To calculate the read coverage for each locus, MHCnvex counts reads mapped to each base pair using Mosdepth (*Pedersen et al. 2018*). To ensure that the analyzed tumor and normal samples have comparable coverage, MHCnvex normalizes raw counts to the total sequencing depth.
2. <u>MHC LR calculation:</u> In order to calculate the LR of the MHC region, coverage of normal samples are used to detect local coverage peaks in MHC loci. Regions in normal samples with coverage peak can provide more reliable measurements as they have more reads mapped to them. By utilizing this approach, the LR calculation becomes independent of the capture panel used for sequencing. If a matched normal sample is not available, MHCnvex can use non-matched normals to find the local peaks. Once the peaks for each gene are identified, adjacent base pairs (default 50 base pairs on each side) are pooled together to form one bin on the gene. Depending on the number of peaks per each gene, multiple bins per gene will form. This approach ensures that there are enough reads per bin to call reliable LR values and reduces noise in the segmentation step. By having the bins and their coverages in the normal and tumor samples, MHCnvex is then able to calculate the log2-ratio values as:

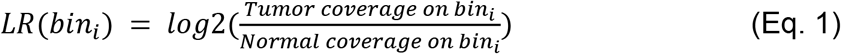
3. <u>MHC BAF calculation:</u> BAF values for the MHC region are calculated by utilizing the coverage of each haplotype determined in the log2-ratio (LR) calculation step. Loci at which the two haplotypes are mismatched are identified as Single Nucleotide Polymorphisms (SNPs). For these mismatch regions, BAF is calculated as:

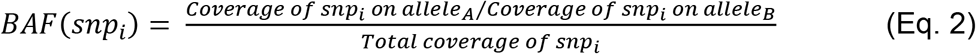
4. <u>Integration:</u> At this step, LR and BAF values for the MHC locus and the regions adjacent to this locus (regions on chromosome 6 before and after MHC locus) are both available. MHCnvex then integrates these values to generate a continuous genomic sequence with their corresponding LR and BAF values. The range of the adjacent region for integration can be set by the user, with a default range being at the whole short arm of chromosome 6.
5. <u>Segmentation:</u> To segment the integrated MHC locus and flanking regions, a custom algorithm is employed based on Circular Binary Segmentation (CBS) to jointly analyze integrated log coverage ratios and BAFs directed toward identifying breakpoints in the tumor’s genome. The algorithm proceeds in two stages: first, independent segmentation of LR and mirrored BAF using CBS with parameters alpha = 0.01 and trim = 0.025, followed by collection of all candidate breakpoints. In the second step, the resulting segmentation track is iteratively refined by merging segments that exhibit similar LR and BAF values, have short lengths, are located in blacklisted regions, and exhibit high variation in coverage among the whole cohort of normal samples. This approach is specifically designed to achieve high sensitivity in detecting breakpoints by initially collecting all possible breakpoints and then selectively removing those with low confidence levels.
6. <u>Model fitting:</u> MHCnvex performs the copy number state inference on the resulting segmented copy-number profiles, Using a custom model most similar to the mathematical formalism of ABSOLUTE (*Carter et al, 2012*) and PureCN (*Riester et al. 2016*). Briefly, the algorithm uses the log-ratios (of bins) and BAFs of individual SNPs and assigns them to their corresponding segments. Next, the likelihood of different possible copy numbers (C, ranging from 0 to 18) are calculated using a likelihood model. The likelihood model is based on the assumption that LR values are distributed normally around a segment described in Eq. 3

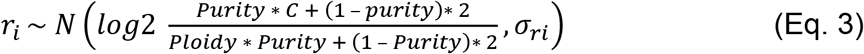

where *σ_ri_* is the average variance of lr values, distributed around a segment. To calculate one likelihood value for each segment, the likelihood of all the bins on a segment are summed up together. Finally, for each segment, the C that yields the highest likelihood is picked. After inferring the C using the LR values for each bin, MHCnvex uses BAFs in the Eq. 4 to determine the expected allele frequency for each SNP:

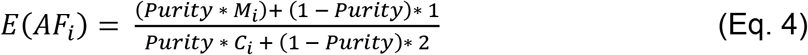 Where *M_i_* is the number of alleles harboring a specific SNP and *C_i_* is the total copy number of the SNP locus. After inferring C through Eq. 3 and both ‘theoretical expected’ (Eq. 4) and observed value for AF (through sequencing) are determined, then a likelihood value can be calculated for different values of *M_i_* according to the equation 5

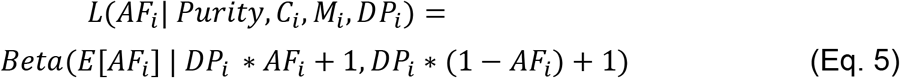

where *DP_i_* is the total depth for *SNP_i_*. By maximizing L in the Eq.5, which is parameterized by M, the multiplicity of each SNP and therefore number of each haplotype can be called.
7. <u>Calling LOH:</u> After having the allele specific copy number for each locus of the genome, MHCnvex can call the LOH for segments which have a minor copy number of 0. MHCnvex also returns the copy number of each HLA haplotype for genes in HLA class 1 and class 2.

**Figure 1:**
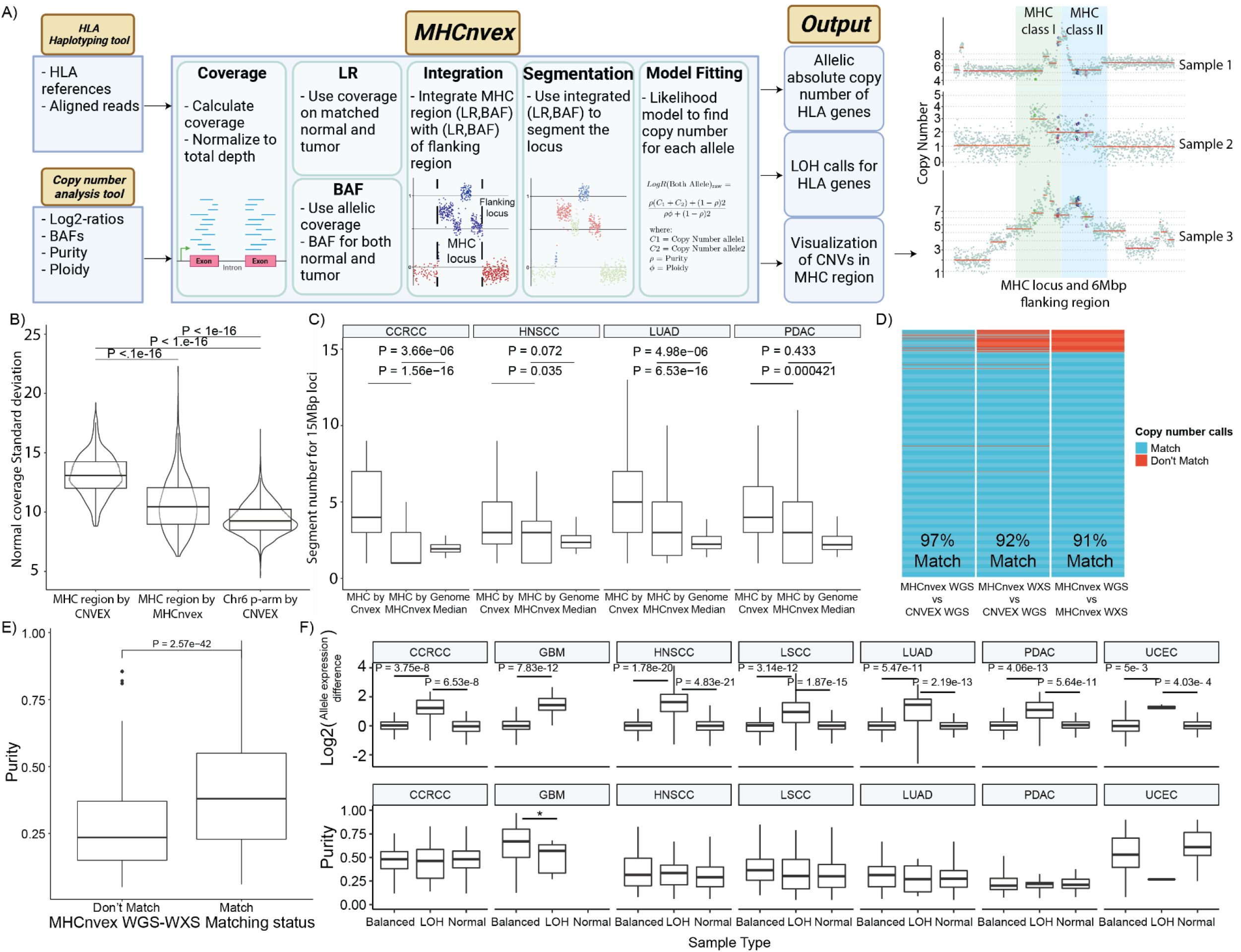
MHCnvex absolute copy number calculation. (A) MHCnvex workflow. MHCnvex accept input from an HLA haplotyping and a CNV analysis tool. Then Works in 7 steps to call CNV and LOH for genes in MHC class-I and class-II. Finally, it outputs allelic copy number of each gene, LOH call for each gene, and a visualization of the loci. 3 examples of the MHC region have been shown (B) Standard deviation (SD) of the normal coverage in the MHC genes. Standard deviation of MHC region calculated by MHCnvex has a significant SD reduction compared to the same value calculated by standard CNV approach CNVEX (C) Number of segments called by MHCnvex versus standard CNV approach CNVEX. MHC region is prone to over segmentation due to polymorphism. MHCnvex shows a significant reduction in the number of segments compared to CNVEX. MHCnvex results are moving toward the median number of segments for every 15MBp tile of the genome (D) MHCnvex WXS versus WGS setting performance. MHCnvex methods shows a high concordance of 91% between the two setting. They also show concordance with the copy number of flanking regions calculated by CNVEX (E) Samples with inconsistence CNV calls in panel D have significantly lower purity compared to the consistent ones (F) Top panel shows the difference in allelic expression of MHC class-I genes. Across all cohorts, samples detected by MHCnvex with MHC class-I LOH have significantly higher difference in expression of their alleles compared to normal samples or tumor samples detected to have balanced allelic number. Bottom panel shows that these different groups across all cohorts have comparable purity and therefor, in this analysis, purity would not affect the difference in expression

### 2.2 WGS analysis

For WGS analysis, first MHCnvex calculates the BAF value of each MHC gene as mentioned above using personalized reference coverages. For the LR calculation in the WGS setting, MHCnvex uses larger bins compared to the WXS setting to ensure there are enough reads per each bin for accurate LR calculation. In the WGS setting, the minimum length of a bin for a reliable estimation of LR value exceeds the length of MHC genes. Hence, in order to get reliable LR values, it is not possible to only use reads mapped to personal references of MHC genes. To overcome this problem, MHCnvex follows the same approach mentioned above to calculate absolute copy numbers for regions adjacent to each gene (LR and BAF values for MHC genes are not used yet). If both sides of a gene had the same copy number value, that value is also assigned to the gene. If they have different copy numbers, then the side with mirrored BAF closest to the mirrored BAF of MHC gene is selected and its copy number is then assigned to the MHC gene. After having the absolute copy number of each gene, the calculated BAF value of the gene itself is used to estimate the allelic copy numbers.

### 2.3 Haplotyping and alignment

In this study, the MHC haplotyping is performed using Hapster, a tool that generates personalized reference sequences for each patient based on their unique MHC haplotypes. Reads are then mapped to these personalized references for both the tumor and its matching normal samples.

### 2.4 Non-MHC CNV calls

CNVEX (*Li et al, 2023*) is used to infer copy number variations for the short arm of chromosome 6. CNVEX is also used to calculate the purity and ploidy of the test data set for validation and results.

### 2.5 Allelic RNA expression differences test

Hapster is used to calculate the RNA expression of each HLA gene’s allele using the personal germline reference. MHCnvex is used to find the copy number profile of those genes. Then to test the difference in the allelic-expression of MHC genes for tumors with LOH versus the MHC balanced (equal allelic copy number) samples, “difference measure” is defined as:

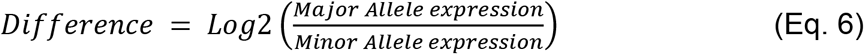

where “major” and “minor” are assigned by MHCnvex for the tumors with MHC LOH, based on their allelic copy number. For all the other cases in balanced and normal groups with equal number of alleles, “major” and “minor” are randomly assigned to the alleles of a gene. The Wilcoxon rank-sum test is used for statistical significance analysis.

## 3. Results

### 3.1 Data acquisition

In this study, whole exome sequencing (WXS), whole genome sequencing (WGS) and polyA RNA sequencing data were obtained for 712 patients from the CPTAC3 pan-cancer cohort. This data was processed using our in-house genomic analysis pipeline, TPO, for variant calling, copy number analysis in genomic data and RNA quantification for RNA sequencing data. This cohort includes 7 different cancers with some of them (LSCC and LUSC) already reported to be enriched with HLA LOH events (*McGranahan et al., 2017*). All the tumor samples have a matching normal except for the GBM cohort.

### 3.2 MHCnvex significantly removes the coverage noise in the MHC region

The accurate calculation of read coverage for each locus is a crucial step in the analysis of copy number variations. Inaccurate coverage estimates can lead to over-segmentation of the genome, resulting in false CNV calls as the algorithm may interpret each segment as a separate CNV event. Therefore, it is imperative to accurately determine the read coverage of MHC locus to ensure the reliability of CNV analysis. In theory, coverage in a normal sample for all the patients should be similar as there are no CNV events, but the polymorphic nature of the MHC region can cause variabilities in the calculated coverage. We evaluated MHCnvex ability to reduce these variabilities compared to conventional CNV analysis tools using a measure of “Coverage variance” in WXS data. Variance in the coverage of non-polymorphic regions of chromosome 6 arm p was used as a baseline since these loci are not prone to errors involved with polymorphism. Our analysis showed that the variance in coverage of the MHC region obtained from the standard approach was significantly higher than the variance expected in coverage of arm p of chromosome 6 (P-value < 1e-16, **Figure 1B**). When the same analysis was conducted using MHCnvex, we observed a significant reduction in variance compared to the standard approach for the MHC region (P-value < 1e-16). However, we still observed a higher variance rate for MHCnvex compared to the baseline which can be explained partly by the possible errors in data processing steps (i.e Haplotyping of MHC region, pooling and library preparation according to the reference genome, etc.) prior to the MHCnvex analysis. Overall, MHCnvex shows significantly less coverage variance among normal samples compared to the same region when coverage is calculated by standard approaches.

### 3.3 MHCnvex significantly reduces the number of segments in the MHC region

Another crucial step in the identification of CNV is the segmentation of loci based on their log-ratio and B allele frequency (BAF) values. However, regions with high levels of variability in coverage values are prone to over-segmentation and subsequently, the generation of faulty CNV calls. This variability is specially increased in the MHC region due to the polymorphism nature of the locus. To evaluate the ability of MHCnvex to avoid over-segmentation, we compared the number of segments produced by MHCnvex to those produced by standard approaches in the same region. A 15Mbp region on chromosome 6, comprising a 3Mbp MHC region and 6Mbp flanking regions on either side, was selected for this analysis. Additionally, we utilized the median number of segments for all other regions of the genome tiled into 15Mbp loci as a baseline.

MHCnvex showed a significant decrease in the number of segments compared to the standard approach (P-value in the figure, **Figure 1C**), which is in the same direction as the baseline number of segments.

### 3.4 MHCnvex shows consistent results using WXS and WGS data

To test the concordance of different methods of CNV calling by MHCnvex, we ran MHCnvex with two different types of input: (i) WGS data, and (ii) WXS data. Copy number calls for flanking regions of MHC genes were also used to assess the concordance of MHCnvex inferred copy numbers with that of flanking regions. In terms of performance, MHCnvex showed a concordance of above 90% with copy numbers of flanking regions for both WGS and WXS settings. MHCnvex (WGS setting) showed a high concordance of 97% with CNVEX (using only WGS data) (**Figure 1D**). Further, comparison of the MHCnvex WXS setting with CNVEX (ran with WGS data) showed a concordance of 92%. Comparison of the results from MHCnvex WXS and MHCnvex WGS settings also showed a high concordance of 91%. Some degree of inconsistency between WXS and WGS can be explained by the higher resolution of WXS setting in detecting focal events which WGS analysis doesn’t have. Next, we compared samples showing inconsistency in CNV calls between the WXS and WGS setting of MHCnvex. Our analysis revealed that samples with mismatch had significantly lower purity (P-Value = 6.62e-05) as shown in **Figure 1E**. This findings are consistent with results from other MHC LOH calling tools, as they have also reported lower accuracy for tumors with low purity (*McGranahan et al., 2017, Pyke et al., 2022*)

### 3.5 MHCnvex LOH calls correlate with changes in RNA expression

Next, we evaluated the accuracy of LOH calls using the RNA allelic-expression differences. Three categories of samples were analyzed: those with LOH detected by MHCnvex, those with equal copy numbers of each haplotype (referred to as “balanced”), and normal samples. Our study specifically focused on MHC class-I genes, as they are highly expressed in all cell types, allowing for the comparison of normal and tumor samples within each cohort. Our results showed that, across all cohorts, the LOH group had a significantly higher difference in allelic RNA expression compared to the other two groups (P-value shown in figure, **Figure 1F**). Additionally, we found that the “balanced” group had similar average differences to the normal group, as expected given their equal allele copy numbers. The average purity of each group within each cohort was also similar, suggesting that purity is unlikely to be a confounding factor in the results (**Figure 1F**).

## 4. Conclusion

In this study, we present MHCnvex, an algorithm for the detection of copy number variation and loss of heterozygosity in the highly polymorphic MHC class I and II regions. MHCnvex is versatile and can be applied to both whole exome sequencing and whole genome sequencing data separately. It can also integrate them (if available) for a better LR and BAF calculations. To further enhance the clinical utility of MHCnvex, it is designed to work in instances without a matched normal for the tumor. One of the key features of MHC is that it takes advantage of the additional information from regions adjacent to the MHC in the process of segmentation, thus removing local biases of the data for the analysis. Our results show that MHCnvex has reduced noise caused by polymorphisms in the region, both at the coverage and segmentation level. Furthermore, we used allelic-expression differences in RNA-seq data to confirm that samples detected to have LOH indeed show significantly higher allelic-expression differences compared to the samples that are detected to have a balanced allelic copy number.

